# A Rapid Inducible RNA Decay system reveals fast mRNA decay in P-bodies

**DOI:** 10.1101/2023.04.26.538452

**Authors:** Lauren A. Blake, Yang Liu, Takanari Inoue, Bin Wu

## Abstract

RNA decay plays a crucial role in regulating mRNA abundance and gene expression. Modulation of RNA degradation is imperative to investigate an RNA’s function. However, information regarding where and how RNA decay occurs remains scarce, partially because existing technologies fail to initiate RNA decay with the spatiotemporal precision or transcript specificity required to capture this stochastic and transient process. Here, we devised a general method that employs inducible tethering of regulatory protein factors to target RNAs and modulate their metabolism. Specifically, we established a Rapid Inducible Decay of RNA (RIDR) technology to degrade target mRNA within minutes. The fast and synchronous induction enabled direct visualization of mRNA decay dynamics in cells with spatiotemporal precision previously unattainable. When applying RIDR to endogenous *ACTB* mRNA, we observed rapid formation and disappearance of RNA granules, which coincided with pre-existing processing bodies (P-bodies). We measured the time-resolved RNA distribution in P-bodies and cytoplasm after induction, and compared different models of P-body function. We determined that mRNAs rapidly decayed in P-bodies upon induction. Additionally, we validated the functional role of P-bodies by knocking down specific a P-body constituent protein and RNA degradation enzyme. This study determined compartmentalized RNA decay kinetics for the first time. Together, RIDR provides a valuable and generalizable tool to study the spatial and temporal RNA metabolism in cells.

## Introduction

RNA is the essential intermediate biomolecule that transmits genetic information encoded in DNA into functional proteins. It is tightly regulated both at its birth (transcription) and death (degradation). RNA degradation is essential to maintain transcript homeostasis or clear defective RNA species. In eukaryotic cells, normal mRNA degradation is initiated by deprotection of its ends with deadenylation or decapping, followed by 5’ → 3’ degradation by XRN1 or the 3’ → 5’ decay by RNA exosome. Defective mRNAs are cleared by RNA quality control pathways; for instance, nonsense-mediated decay (NMD), no-go decay, or non-stop decay^1^. In the NMD pathway, a premature stop codon activates the RNA helicase UPF1, eventually committing the RNA to degradation by recruiting heterodimer SMG5/SMG7 to decay mRNA through the deadenylation/decapping pathway or the endonuclease SMG6 to cleave the mRNA^2^.

RNA-containing membraneless organelles, including nuclear speckles, stress granules (SGs), and processing-bodies (P-bodies), play important roles in RNA metabolism, including splicing, modification, storage, and decay^3, 4^. P-bodies were initially discovered in yeast and were shown to be enriched with RNA decay machineries, but devoid of ribosomes and translation factors^5, 6^. Therefore, it was hypothesized that P-bodies were the sites of RNA decay. However, subsequent studies have challenged this view as RNA degradation still occurs in the absence of visible P-bodies^7, 8^. Recent research suggests that P-bodies might serve as storage sites for mRNAs that can be translated again upon exiting P-bodies^9, 10^. Unlike SGs, P-bodies exist in steady-state physiological conditions, and recruiting RNA to P-bodies typically requires the application of stress, such as amino acid starvation or osmotic stress^11^. During stress, a large variety of mRNAs are recruited to P-bodies, which mediates the stress response and recovery. Currently, the physiological function of P-bodies in unstressed states remains elusive. To address these questions, it is crucial to investigate the localization and degradation of specific RNAs in P-bodies and the cytoplasm separately under physiological conditions.

Conventionally, RNA decay is measured in bulk experiments by harvesting RNA at different time points after inhibiting transcription or pulse labeling of new transcripts^12^. Cells are lysed, and if necessary, fractionated to enrich certain cellular compartments. As a result, the spatial information is lost, and the temporal resolution is limited. Fluorescence imaging tracks the RNA and organelles in real time, which allows for direct visualization of biological events in subcellular compartments with temporal resolution compatible for mRNA decay.

Imaging mRNA at the single molecule level in live cells is crucial for unraveling the mechanism of RNA synthesis, transport, translation, and degradation. Single-cell / single-molecule imaging technology has enabled the direct measurement of transcription dynamics at the single allele level^13^. However, the spatiotemporal dynamics of RNA decay in cells remain poorly understood, due to the transient nature of degradation and the disappearance of signal being frequently confounded by imaging artifacts. While enzymatic degradation of RNA occurs in seconds to minutes, many mRNAs in mammalian cells take hours before they are committed to degradation. The rapid diffusion of mRNAs in cells makes it challenging to track and capture the infrequent and transient decay process. Modulating RNA decay on demand would be instrumental because it can synchronize the transient process^14^. While existing methods, like RNA interference, are convenient to knock down target genes, it takes many hours to exert effects - too slow for studying decay dynamics. Thus, a method for inducing rapid and synchronous decay of RNAs is highly desirable.

In this study, we established a rapid inducible RNA decay system by recruiting an RNA degradation factor on demand, and quantified RNA in subcellular compartments in single molecule resolution. We demonstrated that RIDR can knock down target mRNAs faster than standard small interfering RNA (siRNA). The rapid synchronous decay allowed us to study the function of membraneless organelles, such as P-bodies. By combining RIDR with genetic and pharmacological perturbations, we revealed the functional role of P-bodies in RNA decay.

## Results

### An inducible RNA decay system that is fast and specific

This method is motivated by RNA’s natural propensity to assemble into ribonucleoprotein particles (RNPs). RNA is not a naked polymer of nucleotides; it associates with numerous protein factors, which ultimately determine its fate. The protein composition of RNPs is constantly remodeled during the lifecycle of an mRNA. By tethering specific RNA binding proteins to the target mRNAs, their fate can be artificially influenced. To modulate RNA decay on demand, we implemented a chemically inducible dimerization (CID) system system to control RNA metabolism by fusing a RNA decay factor and a sequence-specific RNA binding protein to a CID pair **(Fig. 1a)**. The CID pair we utilized consisted of the FK506 Binding Protein (FKBP) and the FKBP−Rapamycin Binding domain (FRB) that rapidly dimerize at low concentrations of rapamycin (Rapa)^15^. The FRB/FKBP CID system has been applied to control protein dimerization and many cellular functions before^16, 17^.

**Figure 1:**
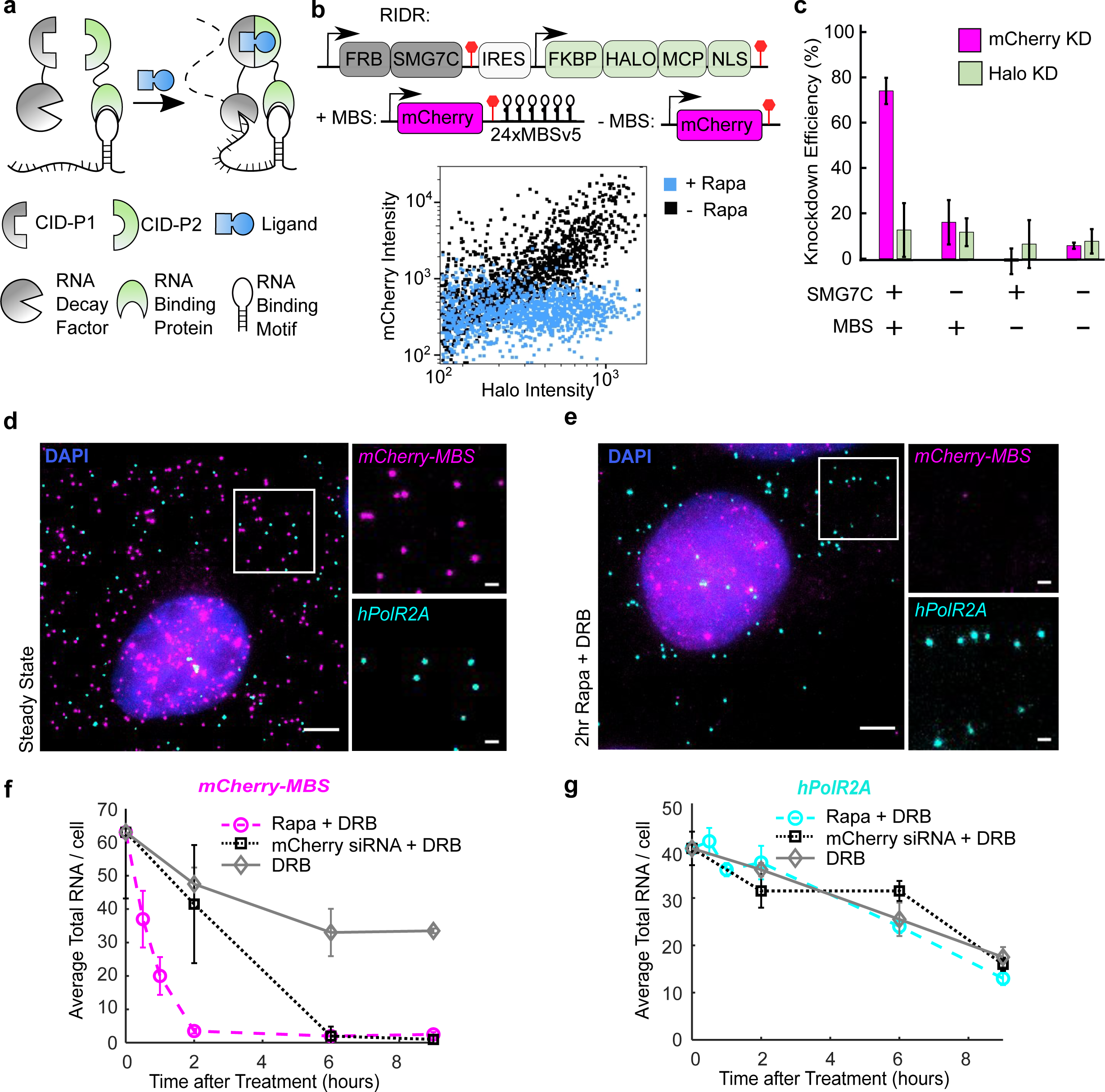
RIDR system is fast, specific, and inducible. **a)** Schematic of a generalized inducible mRNA decay system for inducibly targeting an RNA binding motif with an RNA decay factor. **b)** HEK293T cells were transiently transfected with mCherry-MBS and RIDR constructs. The cells were treated with rapamycin (Cyan) or DMSO control (Black) at the time of transfection. FKBP-HaloTag-tdMCP in the RIDR construct was labelled with JF503-Halo-ligand. The fluorescence of single cells was measured by flow cytometry and presented as the scatter plot. IRES: Internal Ribosome Entry Site. **c)** Knockdown efficiencies of mCherry and HaloTag were quantified from flow cytometry experiments in (b) under conditions listed. The knockdown efficiency is calculated for each condition with respect to itself when Rapa is not added. The HaloTag signal is used as a negative control that does not depend on tethering and Rapa. 1000-3000 HaloTag-positive cells were quantified per condition. Error bars represent the standard deviation across 3 biological replicates. **d-e)** Representative smFISH images of U-2 OS cells stably expressing *mCherry-MBS* and RIDR in steady state condition (**d**) and after 2 hours of rapamycin treatment (**e**). The white box was enlarged on the right. *mCherry-MBS* FISH: magenta; *hPolR2A* FISH: cyan; DAPI: blue. Scale bars: 5 µm for original and 1 µm for zoomed images. **f-g)** Quantification of time-resolved two-color smFISH experiment over 9-hours after induction with Rapa (circle), mCherry siRNA (square), or DMSO (diamond). DRB was added in all experimental conditions to inhibit transcription. The number of transcripts for *mCherry-MBS* (**f**) and *hPolR2A* (**g**) were counted in the same cells for all time points. Rapa + DRB: circles; mCherry siRNA + DRB: squares; DRB alone: diamonds. Error bars represent standard deviation of the means of 2 biological replicates (60-110 cells were quantified per condition in each replicate

The tethered RNA decay factor should meet several criteria. First, it needs to be non-toxic when over-expressed. Second, it should be active only when proximal to the target RNA, with minimal off-target effects. Third, it should efficiently prompt RNA degradation upon tethering. One such promising candidate, SMG7, functions in the nonsense mediated mRNA decay (NMD) pathway. It is one of the last factors recruited in the NMD pathway once an mRNA is committed to decay irreversibly^18^. Previously, it was demonstrated that tethering of the C-terminus of SMG7 (SMG7C) directly to an mRNA decreases its half-life up to 3 times without any upstream NMD factors^19^. Another candidate is the endonuclease in the NMD pathway, SMG6. The catalytic PIN domain (SMG6PIN) can also degrade target RNA when tethered^20^. We compared the efficiency of SMG6 and SMG7 to degrade the target mRNAs when directly tethered. We constructed reporters encoding fluorescent protein mCherry with different numbers of MS2 Binding Sites (MBS) in the 3’ untranslated region (UTR). The plasmids mCherry-nxMBSv5 (where n = 0, 1, 3, 6, 12, 24)^21^ were co-transfected into HEK293T cells with the tandem MS2 Coat Protein (tdMCP) that specifically binds MBS motifs^22^. The expression of mCherry was then measured by flow cytometry **(Fig. S1a)**. We directly fused HaloTag-tdMCP to either SMG7C or SMG6PIN and measured the knockdown efficiency relative to a negative control without any RNA decay factor(**Figs. S1b-c)**^23^. SMG7C can degrade target RNA more efficiently than SMG6PIN, achieving a 68% knockdown efficiency with just 3x MBS, while SMG6PIN required 24 stem loops to achieve similar levels of knockdown **(Fig. S1c).** Therefore, we concentrated on SMG7C in the following experiments.

Next, we employed an inducible tethering system by constructing a bicistronic vector with FRB fused to SMG7C, and FKBP fused to HaloTag-tdMCP, with the latter translated from an internal ribosome entry site (IRES) to facilitate co-expression. We named the construct Rapid Inducible Decay of RNA (RIDR) **(Figs. 1b and S1d).** We then co-transfected RIDR and the mCherry-24xMBS (*mCherry-MBS*) plasmids in HEK293T cells. Flow cytometry results show that mCherry levels decreased 74% upon induction with Rapa **(Figs. 1b-c).** The knockdown is not due to translation repression from Rapa, as it is dependent on tethering of SMG7C by tdMCP **(Fig. 1c)**. Not surprisingly, the addition of more MBS in the target RNA improved the knockdown efficiency **(Fig. S1e).**

To assess the speed of RIDR, we measured the kinetics of the mRNA decay using single-molecule fluorescent in situ hybridization (smFISH) with probes targeting the MBS region. In U-2 Osteosarcoma (U-2 OS) cells stably expressing both the *mCherry-MBS* reporter RNA and the RIDR construct, >90% of *mCherry-MBS* RNA disappeared upon induction for 2 hours **(Figs. 1d-e)**, which was much faster than the gold standard RNA interference using mCherry siRNA treatment **(Fig. 1f).** Importantly, a nontargeting endogenous mRNA, *hPol2RA*, decayed similarly in the Rapa condition, demonstrating the specificity of the RIDR system **(Fig. 1g)**. To avoid confounding with newly synthesized mRNAs, we applied transcription inhibitor 5,6-dichloro-1-beta-D-ribofuranosylbenzimidazole (DRB) in all experiments **(Figs. 1f-g)**.

### RIDR induces rapid decay of endogenously labeled mRNAs

To further examine the effectiveness of RIDR, we applied the tool to endogenous genes tagged with MBS. 24x MBS have previously been knocked into the 3’UTR of mouse β-actin (ACTB-MBS) at the endogenous loci without influence on its function^24^. We stably expressed the RIDR construct in mouse embryonic fibroblasts (MEF) extracted from the mouse, here forth referred to as ACTB-MBS MEF. The expression of *ACTB-MBS* mRNA is significantly higher than the *mCherry-MBS* reporter, but it can be knocked down equally fast: with 95% knockdown at 2 hours post-induction with Rapa. This is significantly faster compared to siRNA treatment, which only achieved 24% knockdown in 2 hours **(Figs. S2a-e).** The decay of endogenous *mPolR2A* mRNA without MBS was not influenced by Rapa induction, demonstrating the specificity of RIDR **(Fig. S2f)**.

### RIDR induces RNA granules formed on pre-existing P-bodies

The rapid and synchronous RNA decay induced by RIDR allows observation of emergent phenomena that are obscured by stochastic decay of single mRNAs. To track the RNA decay in real time, we performed live cell imaging of *ACTB-MBS* mRNAs using FKBP-HaloTag-tdMCP labeled with Janelia Fluorophore (JFX646)^25^. We observed that RNA granules emerged within 5 minutes after induction and disappeared within 1 hour **(Supplemental Movie 1)**. To ascertain the identity of these granules, we simultaneously performed smFISH and immunofluorescence (smFISH-IF) with antibodies against common cytoplasmic RNA granules. First, we observed RIDR did not cause SG formation, as confirmed by staining with G3BP1, a common validated SG marker **(Figs. S3a-b).** Next, we checked if the RNA granules colocalized with P-bodies markers by staining with known P-body markers, decapping enzyme 1A (DCP1a) and DEAD box helicase 6 (DDX6). Though P-bodies are present in steady state conditions **(Fig 2a)**, the ACTB-MBS RNA only colocalized with the DCP1a and DDX6 puncta after induction with Rapa **(Figs. 2b and S3c, respectively).** At 2 hours post-induction, a majority of *ACTB-MBS* RNAs were depleted from both the cytoplasm and the P-bodies **(Fig. 2c)**. In contrast, after treatment with siRNA against ACTB, no bright RNA granules formed that colocalized with P-bodies, even though individual *ACTB-MBS* mRNAs were found within P-bodies occasionally **(Figs. 2d-e)**. The negative control *mPolR2A* mRNA did not accumulate in P-bodies in any treatment **(Fig. 2f).** The P-body number and average intensity did not change within the first hour of treatment. At longer time scales, the number of P-bodies decreased, and their individual average intensity increased **(Figs. 2g-h)**, possibly due to merging of P-bodies at later time points. This is unlikely due to RIDR induction because the P-bodies’ characteristics are similar in all conditions **(Figs. 2g-h)**.

**Figure 2:**
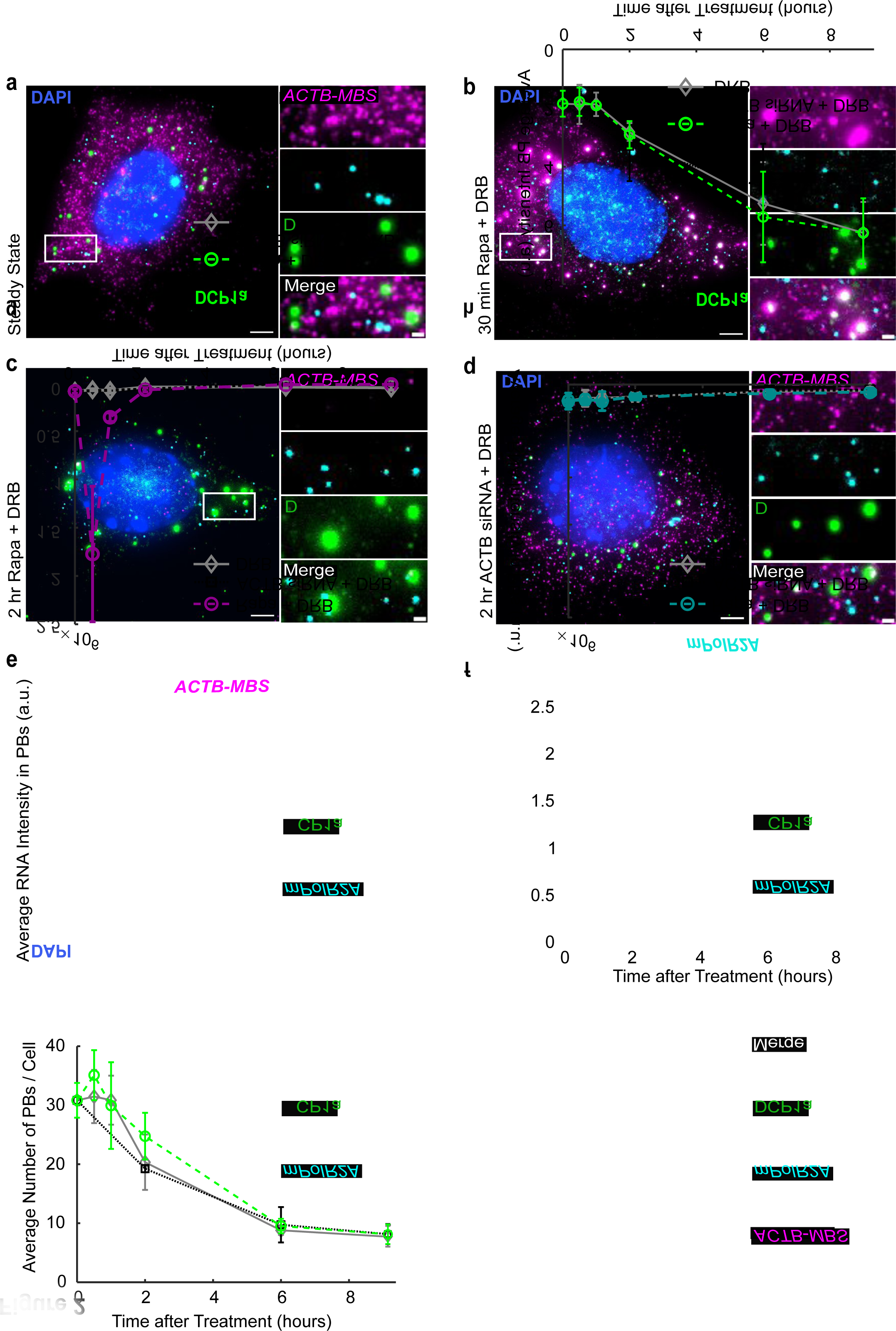
ACTB-MBS transcripts are recruited to P-bodies after RIDR induction. **a-c)** ACTB-MBS MEF cells stably expressing RIDR construct were (**a-c)** induced by Rapa or (**d**) treated with siRNA against ACTB. DRB was added to inhibit transcription at time zero. Cells were fixed at different time points after treatment. smFISH-IF experiments were conducted with FISH probes against *ACTB-MBS* and *mPolR2A*, and antibody against DCP1A. Representative images were shown displaying merged images for FISH and IF channels after **a)** no treatment; **b)** 30-minute Rapa; **c)** 2-hour Rapa; **d)** ACTB siRNA 2-hour treatments. The white box was enlarged on the right. *ACTB-MBS* FISH: magenta; *mPolR2A FISH*: cyan; DCP1a IF: green; DAPI: blue. Scale bars: 5 µm for original images, 1 µm for zoomed images. **e-f)** Quantification of integrated intensities of RNA inside P-bodies after induction, for *ACTB-MBS* (**e**) or *mPolR2A* (**f**) mRNAs. **g-h**) Quantification of P-body number per cell (**g**) and average integrated intensity per P-bodies (**h)**. Rapa + DRB: circles; ACTB siRNA + DRB: squares; DRB alone: diamonds. Error bars represent the standard deviation of the means of 4 replicates (150-284 cells were quantified per condition)

### The effect of translation inhibition on induced RNA decay

Previous studies have shown that translation and RNA decay are intimately coupled^26, 27^. On one hand, ribosomes may compete with decay machinery for access to mRNA and protect mRNA from degradation^28, 29^. On the other hand, it was recently shown that translation may increase mRNA decay rates^30^. To test how translation influences the induced rapid RNA decay, we performed the RIDR experiment in the presence of various translation inhibitors. Cycloheximide (CHX) binds to the ribosomal E-site and inhibits elongation^31^. At high concentration, it freezes ribosomes on transcripts and disperses P-bodies^6^. Indeed, upon CHX treatment, P-bodies disappeared, and RNA granules no longer formed after induction **(Figs. S4a-b)**. The decay of mRNA was also significantly reduced (**Figs. S4c-d**). Another translation inhibitor, puromycin, releases nascent peptides and ribosomes from mRNAs. When treated with puromycin, *ACTB-MBS* mRNAs were recruited to P-bodies after induction and the decay was as efficient as the control (**Figs. S4a-d**) This is consistent with the model that loaded ribosomes inhibit rapid decay of mRNA.

### P-bodies provide a kinetic advantage to RNA decay

In the last section, we demonstrated that after induction, *ACTB-MBS* RNAs were rapidly recruited to P-bodies and disappeared-potentially degraded inside the P-bodies. However, P-bodies are not required for RNA degradation^7, 8^. It is unclear what the function is for recruiting RNA to P-bodies during RNA decay. It is plausible that P-bodies offer a kinetic advantage to decay of RNA due to locally concentrated decay factors^7^. Yet, previously it was difficult to visualize mRNA decay in P-bodies due to their stochastic recruitment of RNA and fast decay speed. The synchronous recruitment induced by RIDR amplified the RNA signal in P-bodies, allowing us to quantify the compartment specific decay kinetics. We developed a kinetic model to describe the RNA trafficking into P-bodies and the subsequent decay dynamics **(Figs. 3a-b and Methods)**. In the most general form, RNAs are recruited into and released from P-bodies with rates *k_R_* and *k_L_*, respectively, and the RNA decay rates in P-bodies and cytoplasm are described by *k_PB_* and *k_CT_*, respectively. To measure these rates, we performed a time-resolved smFISH-IF experiment by fixing cells at different time points after induction. We quantified the number of single mRNAs in the cytoplasm and the integrated intensity of RNA granules in the P-bodies separately (Methods). We normalized the RNA granule intensity into RNA counts by dividing it with the single mRNA intensity (Methods). As a result, we obtained the numbers of mRNAs in the cytoplasm and P-bodies as a function of time. In principle, we could fit the two curves to determine all four rate constants. However, there were not enough features in the curves to unambiguously determine all parameters **(Fig. S5a)**. Therefore, we considered three mechanistically interesting special scenarios with only three independent parameters. First, the P-bodies only functioned as a storage site and there was no decay inside (*k_PB_* = 0). Second, the decay rates in the P-body and in cytoplasm were the same (*k_PB_* = *k_CT_*). Third, once mRNAs entered P-bodies, the rate of leaving was negligible (*k_L_* = 0). The first two models failed to describe the data **(Figs. 3c-d,f)**. The third model described the data equally well with the full model **(Fig. 3e-f, S5a-b).** According to the principle of Occam’s razor^32^, a three-parameter model (Assumption III, *k_L_* = 0) is sufficient. Importantly, the fitting revealed that the decay rate in P-bodies was significantly higher than that in the cytoplasm **(Fig. 3g)**. Taken together, this data suggests that RNA decay can occur in P-bodies and that P-bodies offer a kinetic advantage for RNA decay.

**Figure 3:**
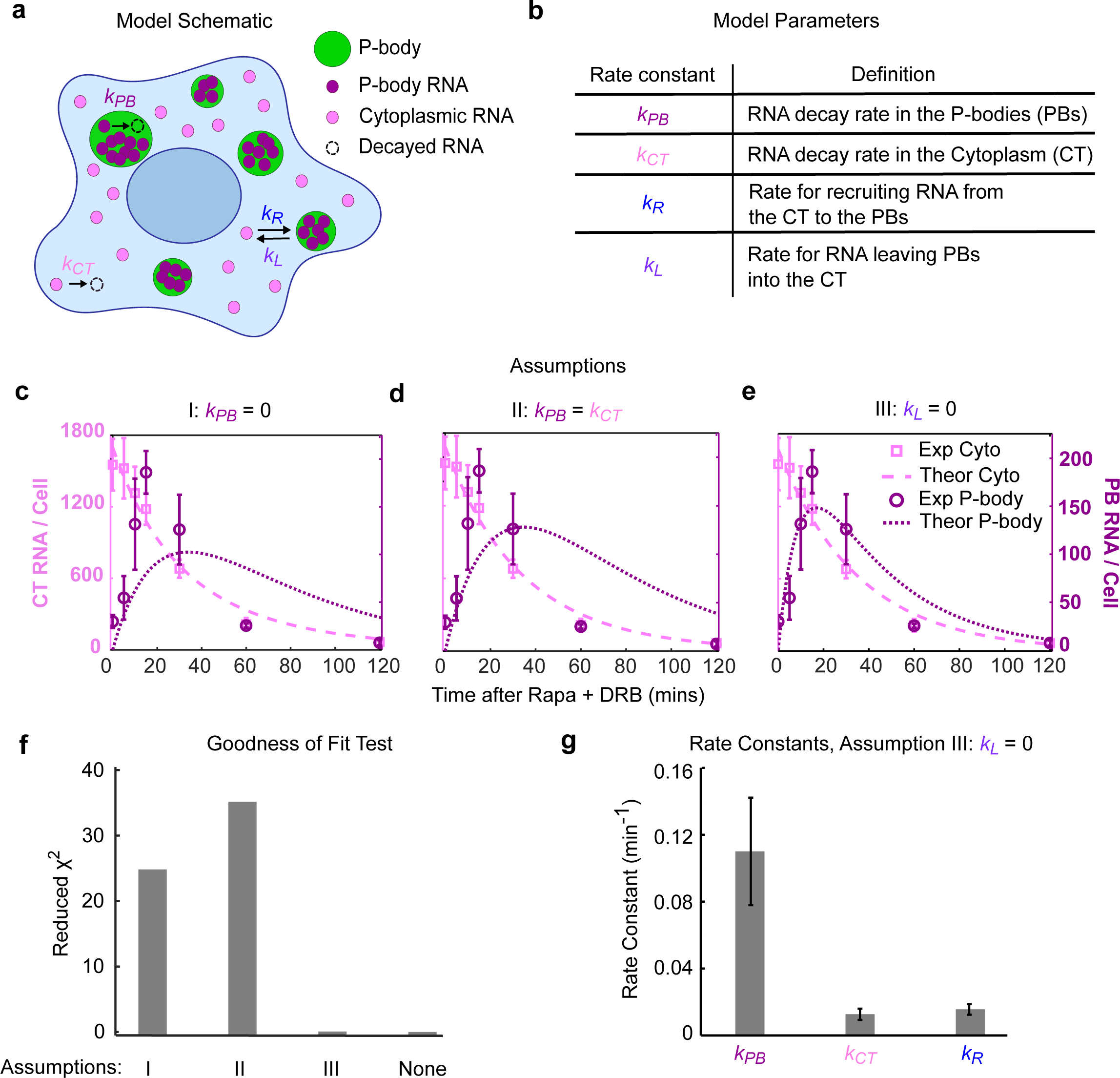
Kinetic Modeling of induced RNA decay in P-bodies and Cytoplasm. **a)** Schematic of the mathematical model depicting RNA decay, recruitment into and release from P-bodies. The details are described in main text and Supplementary Theory**. b)** Table with definitions of each kinetic rate constant. **c-e)** Fitting results under different assumptions **c)** I, no decay in P-bodies:, *k_PB_* = 0; **d)** II, decay in P-body and cytoplasm are the same: *k_PB_* = *k_CT_*; or **e)** III, RNAs recruited into P-body do not leave: *k_L_* = 0. RNA counts in the P-body: dark magenta; RNA counts in the cytoplasm: light magenta; Experimental data: symbols; Theoretical fit: lines. Error bars represent standard deviation of the means of 4 replicates. **f)** Reduced χ^2^ values indicating goodness of fit for models in (c-e), and with full model **(Fig. S5b-c).** Lower values indicate better fitting. **g)** Model parameters determined from fitting with Assumption III: *k_L_* = 0. Error bars indicate the standard deviation of the fitted parameters across 4 replicates (150-284 cells were quantified per condition).

### Fast RNA decay occurs inside the P-bodies

Using mathematical modeling, we showed that RNA decay could occur in P-bodies with a faster rate than in the cytoplasm. To validate this model further, we perturbed the system by depleting key P-body constituent protein or decay enzymes to observe the difference in induced RNA decay kinetics.

Although we observed RNA granules rapidly formed in P-bodies and quickly dissolved, it was still possible that RNA was just processed there and then released into the cytoplasm for decay. We demonstrated that this model **(Fig. 3c, Assumption I**) could not describe the decay kinetics. To convey this directly, we used siRNA to knock down XRN1 (**Figs. S6a-b)**, the major 5’ → 3’ RNA exonuclease which is also enriched in P-bodies^33^. In XRN1 depleted cells, *ACTB-MBS* mRNAs were enriched significantly more in P-bodies and the RNA granules persisted much longer compared to a scrambled non-targeting siRNA control (NC siRNA) treatment **(Figs. 4b and S6c)**. Control mRNAs *mGAPDH* and *mPolR2A* were not recruited nor retained in P-bodies after XRN1 or NC siRNA treatment **(Figs. 4c and S6d, respectively)**. This is consistent with the Assumption III that induced mRNA are recruited into P-bodies and decayed there, as reduced XRN1 levels prolonged the residence time and increased the decaying mRNA levels in the P-bodies.

**Figure 4:**
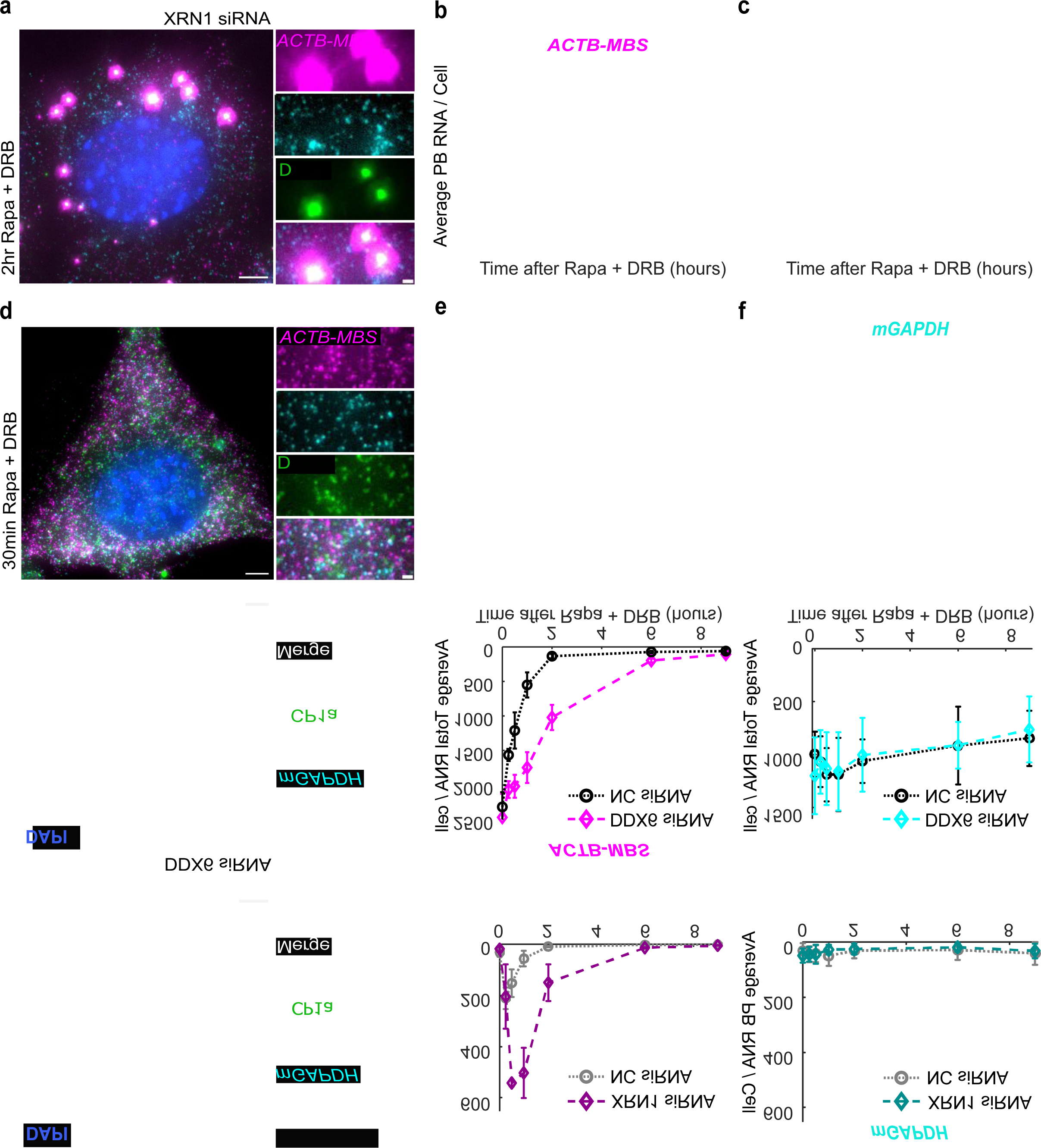
P-bodies are the site of fast RNA decay. ACT-MBS MEF cells were treated with siRNA against (**a-c)** XRN1, (**d-f**) DDX6, or **(a-f)** scrambled siRNA (NC) for 72 hours. Cells were treated with Rapa and DRB, and fixed at different time points. smFISH-IF experiments were conducted with probes against *ACTB-MBS* and *mGAPDH*, and antibodies against DCP1a. **a**) Representative image at 2 hours post induction for XRN1 siRNA treated cells. The white box was enlarged on the right. *ACTB-MBS* FISH: magenta; *mGAPDH* FISH: cyan; DCP1a IF: green; DAPI: blue. Quantification of *ACTB-MBS* (**b**) or *mGAPDH* (**c**) mRNAs in P-bodies over 9-hour time course after induction. XRN1 siRNA: diamonds; NC siRNA: circles. **d)** Representative image for DDX6 siRNA treated cell at 30 minutes post-induction. The white box was enlarged on the right. *ACTB-MBS* FISH: magenta; *mGAPDH* FISH cyan; DCP1a IF: green; DAPI: blue. There is no visible P-body under the DDX6 knockdown condition. **e-f)** Quantification of total *ACTB-MBS* **(e)** and *mGAPDH* (**f**) mRNA levels over 9-hour time course after induction. DDX6 siRNA: diamonds; NC siRNA: circles. Scale bars: 5 µm for original images, 1 µm for zoomed images. Error bars represent the standard deviation of the means of 4 replicates (203-450 cells were quantified per condition).

DDX6, a DEAD box RNA helicase, is essential for P-body formation^34^. When DDX6 was knocked down with siRNA (**Figs. S6c-d)**, no visible P-bodies were observed **(Fig. 4d)**. As a result, no RNA granules appeared after RIDR treatment, as expected **(Fig. 4d)**. *ACTB-MBS* mRNAs still decay without P-bodies, but the rate of decay was slower compared to NC siRNA treatment **(Fig. 4e)**. Importantly, the non-targeting mRNA *mGAPDH* and *mPolR2A* decayed with the same rate when DDX6 was knocked down **(Figs. 4f and S6h)**, indicating that the loss of DDX6 itself does not slow down general RNA decay. Taken together, this data suggests that P-bodies were required to achieve rapid induced RNA decay.

### RNA decay in P-body is sensitive to stress

In the next set of experiments, we aimed to visualize single mRNAs’ recruitment to and decay in P-bodies. There have been controversies about the exact role of P-bodies in RNA metabolism. We observed that mRNAs were recruited to P-bodies and rapidly decayed there. It was puzzling for us that different laboratories have drawn quite contradictory conclusions. During an attempt to capture the RNA dynamics in P-bodies via live-cell imaging, we gained some clues.

We visualized *ACTB-MBS* mRNAs using FKBP-HaloTag-tdMCP^22, 35^ in live-cell imaging. To label the P-bodies, we stably expressed DDX6-eGFP at low concentration in the ACTB-MBS MEF cells. Surprisingly, when we employed the laser power normally used to track single mRNAs, we found that the mRNA recruited into P-bodies and persisted as long as two hours after Rapa induction **(Fig. 5a and Supplemental Movie 2).** To verify that the microscope stage top incubation environment did not perturb the system, we used minimal excitation required for visualizing RNA granules. Indeed, at this condition, we observed the recruitment and dissolution of RNA granules in the P-bodies within one hour, just like the fixed cell experiments where no laser excitation was used **(Fig. 5b and Supplemental Movie 3).**

**Figure 5:**
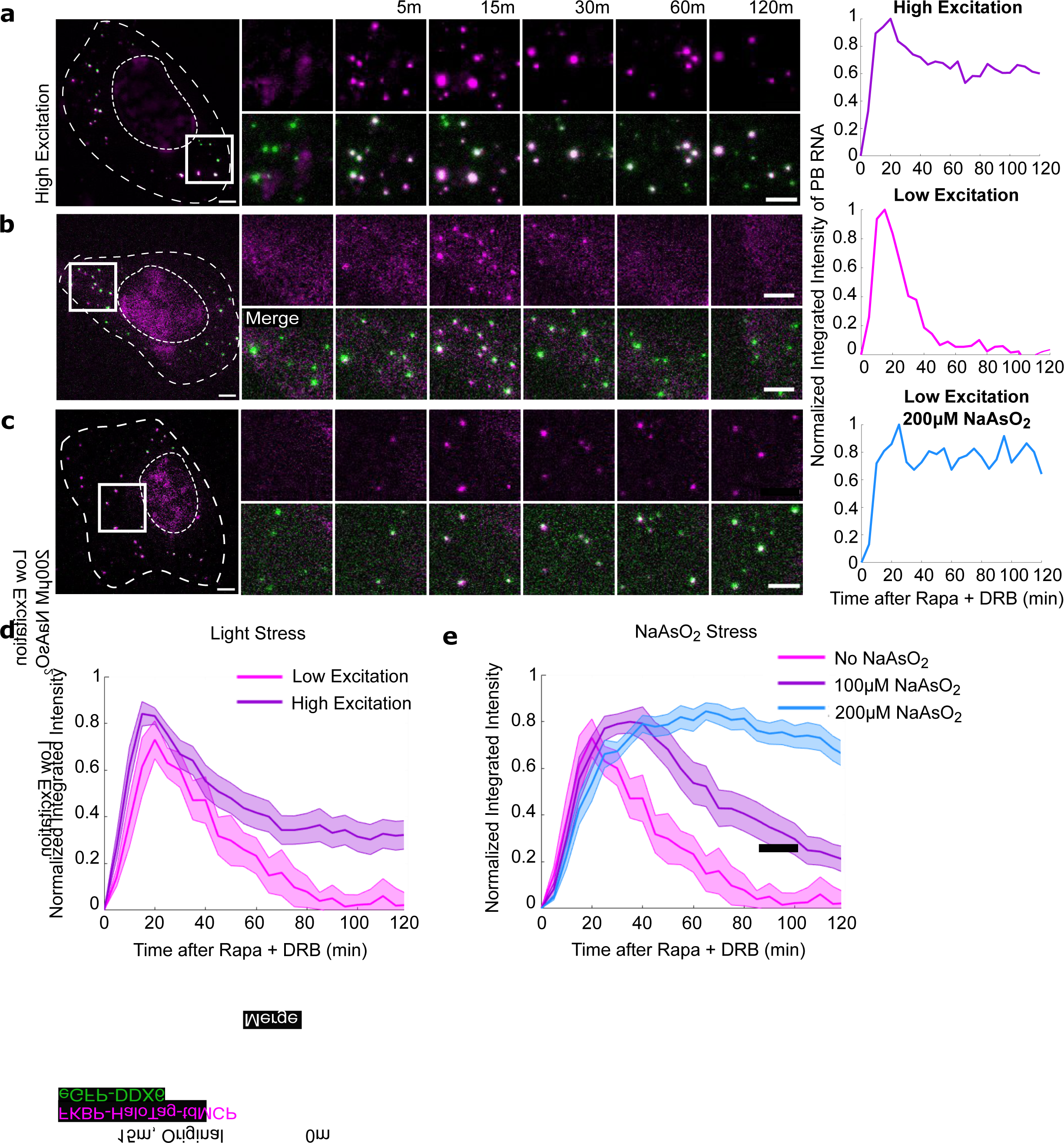
RNA fate in P-body is influenced by stress. Live cell imaging experiments were performed to track P-bodies (DDX6-eGFP, green) and *ACTB-MBS* (FKBP-HaloTag-tdMCP, magenta) after induction. (**a**-**c**) Representative movie montages under different conditions: **a)** High excitation laser power for single mRNA tracking was used **(Supplemental Movie 2); b**) minimal laser power sufficient to observe RNA granules colocalized with P-bodies (**Supplemental Movie 3)**; **c**) minimal laser power and cells were pretreated with 200 µM Sodium Arsenite for 30 minutes (**Supplemental Movie 4).** The normalized intensity traces of RNA granule intensity of movies (a-c) were shown on the right. Scale bars: 5 µm for both original and zoomed images. **d)** Average intensities of RNA granules colocalized with P-bodies under low (magenta) and high (purple) excitation power (number of cells: 8, 14, respectively) **e)** Average intensities of RNA granules colocalized with P-bodies under low excitation when cells were pretreated for 30 minutes with 0 µM (magenta), 100 µM (purple), and 200 µM (blue) NaAsO2 (number of cells: 8, 10, 12, respectively). All intensity traces were normalized in each cell before averaging. Shaded error bars represent standard error.

We hypothesized that the high laser power used for single mRNA imaging created reactive oxygen species and oxidative stress to the cells, hence perturbing the function of P-bodies and shifting the RNA decay kinetics. To verify this hypothesis, we applied artificial oxidative stress to the cells by pre-treating the samples with varying concentrations of Sodium Arsenite (NaAsO_2_) for 30 minutes. We imaged the pre-stressed cells after induction at low excitation conditions **(Fig. 5c and Supplemental Movie 4).** With increasing concentration of NaAsO_2_, mRNAs recruited to P-bodies persisted longer **(Figs. 5c,e)**. This suggests that RNA decay dynamics inside P-bodies are tunable, and the function of P-body decay is context-dependent.

## Discussion

In this study, we developed a genetically encoded inducible RNA decay system that is fast, specific, and modular. We demonstrated its utility for decaying exogenous as well as endogenous transcripts. RIDR is significantly faster than siRNA, reducing the RNA’s half-life from 2-3 hours using siRNA to ∼30 minutes using RIDR. After induction, we observed that endogenous *ACTB-MBS* mRNAs were recruited to P-bodies. We examined the compartmentalized RNA decay with mathematical modeling and genetic perturbation. We concluded that RNA decay occurs inside P-bodies and that P-bodies confer a faster decay rate. Surprisingly, we found that the functional role of P-bodies is modulated by cellular stress. Coupling fluorescence imaging with the synchronous induction of RNA decay, RIDR enables investigation of spatiotemporal RNA decay dynamics previously unattainable. The rapid and synchronous decay is important, as it amplifies the RNA signals recruited into P-bodies, allowing us to determine the distinct compartment-specific RNA decay kinetics.

By measuring the recruitment and disappearance of RNA in P-bodies, this study revealed their functional role. First, we showed that RNA can be degraded in P-bodies. The alternative model that there is no decay in P-bodies cannot fit the experimental data **(Fig. 3c, Assumption I)**. Moreover, when the 5’->3’ exonuclease XRN1 was knocked down, the mRNA enriched in P-bodies increased nearly threefold and required more time to disappear, further supporting the model. These findings challenge the notion that P-bodies are only sites for RNA storage. Second, this study suggests P-bodies provide a kinetic advantage for rapid RNA decay. The rate of decay in P-bodies was faster than in the cytoplasm **(Fig. 3g)**. When P-bodies were disrupted by knocking down the essential constituent protein DDX6, no RNA granules appeared, and the rate of total RNA decay was significantly reduced. While DDX6 itself may influence the RNA decay, we showed that it was not the case because the control *mGAPDH* and *mPolR2A* mRNAs were not influenced by DDX6 knockdown. Rapid decay is required for certain transcripts regulating cell cycle, apoptosis, and embryonic development^36, 37^. It would be interesting to investigate whether P-bodies play a role in regulating the decay of these transcripts. Third, we showed that the P-body function was tunable and modulated by cellular stress. This tunability may underly the variability in the literature regarding the role of P-bodies, as we explain next. There have been controversies about the exact role of P-bodies in RNA metabolism. P-bodies were originally proposed as the location of RNA decay because of the enrichment of RNA decay factors^33^. Later, it was found that RNA decay does not require the presence of visible P-bodies^7, 8^ and P-bodies may have dual roles of RNA storage and decay^38^. Even further, it was proposed that P-bodies function mostly as a place for storing mRNA and that there is no decay in P-bodies at all^39^. In previous single molecule studies, mRNAs were observed entering the P-bodies, then remaining there or leaving without decaying^11, 40^. To obtain a high signal to noise ratio necessary for visualization of single RNAs, one must typically use high intensity imaging conditions. In this study, the RNA recruitment to P-bodies was synchronized, and therefore amplified the signal in the P-bodies. This amplified signal of the RNA enabled the use of a mild, low intensity imaging condition, which revealed a different phenotype than when we employed higher intensity imaging conditions in an attempt to capture single RNA dynamics. The ability to tune the RNA decay dynamics in the P-bodies under varying levels of stress highlights the context-dependence of RNA fate in P-bodies^41^.

The rapid and synchronous RNA decay of highly expressed endogenous transcripts rendered an amplified effect of RNA recruitment to P-bodies upon RNA decay induction. Compared to RNAi, RIDR is significantly faster, therefore, the same observations and kinetic modeling could not be extracted with siRNA treatment. Though we did occasionally observe RNA colocalized with P-bodies after siRNA treatment, it didn’t accumulate to the degree observed with RIDR. This could be due to the slower kinetics of RNAi compared to RIDR, or perhaps the pathway for RNAi does not result in RNA being recruited to P-bodies. More research must be done to determine the factors that determine which RNAs are brought to P-bodies, and whether they will be stored or decayed upon localization. Though we observed SMG7C tethering caused mRNAs to enter P-bodies for decay, it is possible that the fate of decay in P-bodies is also dependent on the context of the SMG7C tethering.

There are some limitations in the current implementation in RIDR. First, we used the MBS/tdMCP system to tether one CID component to RNA. Though CRISPR technology has revolutionized gene editing^42^, knocking in long tags is still cumbersome. A future improvement would be to target unmodified endogenous RNA with programmable RNA binding proteins like rCas9^43^ or catalytically dead CRISPR-Cas13^44–46^. However, we have shown that multiple binding sites are required to achieve efficient knock down with SMG7. Signal amplification would be required to target nonrepetitive mRNAs^47, 48^. Second, the FKBP/FRB CID system requires rapamycin, an inhibitor for mTor signaling that regulates mRNA translation. We have used low rapamycin concentrations such that translation was not significantly affected **(Fig 1c)**, so the rapid response of RNA decay should be independent of the mTOR signaling. In the future, other inducible dimerization systems, such as Giberellin^49^ or light-induced dimerizers^50, 51^ can be used to overcome this limitation.

There are previous efforts to manipulate mRNA metabolism in an inducible manner. While sequestering RNA in artificial clusters can offer translational control, it cannot be used to study the RNA’s metabolism in physiological contexts^52^. Uncaging methods are another way to inducibly influence RNA functions^53, 54^. RIDR provides similar speed and robust tethering efficiency as optogenetic methods developed by Liu and colleagues^55^. The RIDR platform described here is modular, with individual components readily swappable. One can tether other decay factors or RNA regulation factors inducibly to control the mRNA metabolism on demand. By exerting precise spatial and temporal control, the inducible RNA tethering strategy can be used to probe the elusive transient processes that are difficult to study.

## Materials and Methods

### Materials availability

Reagents and materials produced in this study are available from Bin Wu pending a completed Materials Transfer Agreement. Constructs will be made available on Addgene.

### Plasmid construction

The mCherry-24xMBS plasmid was described by Wang and colleagues^56^. To clone the mCherry-nxMBSv5 reporters (where n = 0, 1, 3, 6, 12), we used restriction digestion to remove mCherry-24xMBSv5 from the phage-ubc backbone, then used PCR to amplify mCherry plus the number of MBSv5 desired and ligated the mCherry plus nxMBSv5 stem loops into the original backbone. Because MBSv5 is non-repetitive, the PCR of the stem loops was possible.

For direct tethering of RNA decay factor, we cloned SMG7C-Halo-tdMCP-NLS-HA and SMG6PIN-Halo-tdMCP using Gibson assembly. For the inducible RNA decay factor tethering assays focused on SMG7C, we constructed FRB-SMG7C-IRES-FKBP-Halotag-tdMCP-NLS-HA (RIDR) via 4-part Gibson assembly to clone the RIDR construct into a phage lentiviral backbone with a ubc promoter. An FRB-SMG7C Geneblock was ordered to simplify the cloning. To create FRB-BFP-IRES-FKBP-Halotag-tdMCP-NLS-HA (-SMG7C negative control), SMG7C was replaced by BFP using restriction digestion cloning. Prior to stable cell line integration into U-2 OS or ACTB-MBS MEF cells, the RIDR construct was extracted by PCR and cloned into a Tet-On 3G backbone for Dox-inducibility. The Tet-On 3G backbone was a gift from Sergi Regot’s lab. The inducible expression of RIDR prevents the gene from being lost during long term cell culture.

For live-cell imaging of P-bodies, the phage–UbiC-tagRFP-DDX6 plasmid was ordered from AddGene (#119947) and tagRFP was replaced by eGFP prior to stable integration into the ACTB-MBS MEF cell lines.

### Stable cell line generation

Lentiviral particles were generated by transfecting low-passage HEK293T cells with either FRB-SMG7C-IRES-FKBP-Halo-tdMCP, mCherry-24xMBSv5, or eGFP-DDX6 plasmids along with Generation II viral packaging accessory plasmids [REF?]. Plasmid transfections were performed using polyethyleneimine (PEI). 48 hours following transfection, the viral supernatant was collected, spun down to remove cellular contents, and filtered through a 0.45 µm PVDF filter (Millipore SLHV013SL). The filtered supernatant was applied directly to U-2 OS cells (American Type Culture Collection HTB-96). Viral transduction was performed sequentially by first infecting U-2 OS cells with mCherry-24xMBSv5 and performing fluorescence activated cell sorting (FACS) for mCherry positive cells. This positive population was then infected in the same manner with FRB-SMG7C-IRES-FKBP-Halo-tdMCP and sorted for high–expression cells.

Immortalized MEF cells with 24x MBS at the endogenous ACTB locus (ACTB-MBS MEF) were a gift from Robert Singer’s lab. The ACTB-MBS MEF cells were stably integrated with the dox-inducible expression of FRB-SMG7C-IRES-FKBP-Halo-tdMCP in the Tet-On 3G system and sorted for HaloTag expression using flow cytometry. ACTB-MBS MEF cells used for live-cell imaging were infected in the same manner as above with DDX6-eGFP plasmid for stable integration, and then sorted for GFP expression using flow cytometry.

### Cell culture and transfection

U-2 OS (American Type Culture Collection HTB-96), HEK293T (American Type Culture Collection CRL-1573), and ACTB-MBS MEF cells were grown in DMEM (Corning, 10-013-CV) supplemented with 10% (v/v) FBS (Millipore Sigma, F4135-500ML), 100 U/ml penicillin, and 100 µg/ml streptomycin (Millipore Sigma, P0781) and maintained at 37 °C and 5% CO2. Cells were passaged every 2-3 days once they reached ∼75% confluency. Cells were tested monthly for mycoplasma infection and were always negative.

XtremeGeneHP was used to transfect plasmids into HEK293T cells for use in flow cytometry. For 24-well dishes, each well received 250 ng total plasmid DNA. For flow cytometry experiments, 50 ng of plasmid DNA for the reporter mRNA and 200 ng plasmid DNA containing the RNA decay factor were mixed with serum-free DMEM to a final volume of in 25µL. For each well of a 24-well dish, 1 µL XtremeGeneHP was combined with 24µL of serum-free DMEM and incubated in a MasterMix for 5 minutes at room temperature. After incubation, 25µL of plasmid DNA mix and 25µL of incubated XtremeGeneHP mixture were combined by gentle pipetting, then incubated together for 15 minutes at room temperature. After the second incubation, 50µL XtremeGeneHP and plasmid DNA mixture was added dropwise to each corresponding well. Immediately following transfection, Rapa or DMSO control were added to the cells. Cells were transfected overnight, then prepared for flow cytometry the next morning.

For RIDR vs RNAi benchmarking experiments **(Figs. 1-2)** cells were prepared in the manner described in the RIDR kinetics section. Lipofectamine® RNAiMAX (Thermo Fisher) was used to transfect siRNAs. ON-TARGET pooled siRNAs of ACTB (IDT mm.Ri.Actb.13.1-3), custom designed mCherry siRNAs or OFF-TARGET IDT negative DsiRNAs (NC) were used. For each well of a 24-well dish, a mix of 5 pmol (1 µL of 5 µM pooled siRNA) + 24 µL OptiMem was combined with a mixture of 1.5 µL RNAiMAX + 23.5 µL OptiMem for a total of 50 µL per well. The siRNA / OptiMem / RNAiMAX mixture was mixed gently, incubated for 5 minutes at room temperature, then added dropwise into each well of 24-well. DRB was also added at the time of siRNA treatment, at a final concentration of 100µM. DRB was also added at the time of siRNA treatment, at a final concentration of 100µM.

For DDX6 and XRN1 knockdown experiments **(Fig 4)** Lipofectamine® RNAiMAX was also used to transfect siRNAs. ON-TARGET pooled siRNAs of DDX6 (IDT mm.Ri.Ddx6.13.1-3), XRN1 (IDT mm.Ri.Xrn1.13.1-3) and OFF-TARGET IDT negative DsiRNAs were used. On the evening of Day 1, 100,000 ACTB-MBS MEF cells were plated per well of a 6 well plate. Cells were incubated overnight. On the morning of Day 2, for each well of a 6-well dish, the first RNAiMAX transfection was performed with 7.5 µL RNAiMAX + 67.5 µL OptiMem combined and added to a mixture of 25 pmol siRNA (5 µL of 5 µM pooled siRNA) + 70 µL OptiMem, for a total of 150 µL. The siRNA + OptiMem was mixed gently, incubated for 5 mins at room temp, then added dropwise into the 6-well dish. The cells were incubated for 24 hours. On the morning of Day 3, 15,000 cells were replated onto 12 mm fibronectin-coated coverslips (Electron Microscopy Science, 72290-03) in a 24 well. At the end of Day 3, the second transfection was performed. For each well of 24-well dish, a mix of 5 pmol (1 µL of 5 µM pooled siRNA) + 24 µL OptiMem was combined with a mixture of 1.5 µL RNAiMAX + 23.5 µL OptiMem for a total of 50 µL per well. The siRNA and OptiMem were mixed gently, incubated for 5 minutes at room temperature, then added dropwise into each well of 24-well with coverslips. At the end of Day 4, the medium was replaced. To induce expression of RIDR construct, 1 µg/mL Doxycycline was added to the medium and incubated overnight. On the morning of Day 5, ∼72 hours after initial transfection, the cells were ready for RIDR treatment. siRNA sequences can be found in **Supplemental Table 1**.

### Flow cytometry

HEK293T cells were plated onto a 24-well with 25,000 cells per well, incubated for 24 hours. The cells were transiently transfected with 250 ng total plasmid using the transfection reagent XtremeGeneHP according to the manufacture’s instruction. 100nM rapamycin or DMSO were applied to the cell immediately after transfection. After 14-16 hours, the cells were labelled with 10nM JF503-Halo-Ligand for 1 hour, washed for 30 minutes, trypsinized, resuspended into complete DMEM, then filtered through a cell strainer (Corning 352235). Flow cytometry data was collected on a Thermo Attune NxT flow cytometer. To calculate knockdown efficiency, the JF503-positive cells were gated **(Fig. S1a)** and the geometric mean of fluorescence intensities for mCherry and JF503 channels were calculated in the FlowJo software individually, and compared to the respective control conditions (-Rapa, -SMG7C).

### RIDR time course experiments

For RIDR time course experiments involving U-2 OS cells expressing mCherry-24xMBSv5 **(Fig. 1)**, 50,000 cells were plated the night before on 12 mm coverslips (Electron Microscopy Service, 72290-03). For fixed-cell RIDR time course experiments involving ACTB-MBS MEF cells **(Fig. 2)**, 25,000 cells were plated on fibronectin-coated 12 mm coverslips the evening before a time course experiment. Plating procedures for RIDR kinetics experiments after siRNA treatment **(Fig 4)** or live-cell imaging **(Fig 5)** are explained in the Cell Culture and Transfection section.

For all time course experiments **(Figs.1-5)** cells were treated overnight with 1µg/mL Doxycycline to induce expression of the RIDR construct. The next morning, cells were labelled with JF646 Halo-ligand at the start of the time course experiment prior to the addition of Rapa or DRB Rapamycin powder (LC Laboratories, R-5000-100MG) was dissolved in DMSO for a final stock concentration of 10mM. Rapa aliquots were stored at –20°C for long term storage. Prior to a RIDR experiment, a fresh Rapa aliquot would be used and diluted further to 100µM in DMSO. Transcription inhibitor 5,6-dichloro-1-beta-D-ribofuranosylbenzimidazole (DRB) powder (Millipore Sigma, D1916-50MG) was dissolved in DMSO for a final stock concentration of 100mM. DRB aliquots were stored at –20°C for long term storage and fresh aliquots were used for each experiment.

Both Rapa and DRB were further diluted 1000x to reach their final concentration in the cell culture medium. To ensure adequate dispersion of the drug(s), the appropriate amount of Rapa and/or DRB was added to a fresh microcentrifuge tube, ∼200µL of media was removed from the well or dish where the drugs were to be added, the drug(s) were resuspended in this medium by pipetting, then added back into the well or dish drop-wise. Final concentrations in the wells or dishes for Rapa and DRB were 100nM and 100µM, respectively. Steady State condition was not treated with Rapa or DRB. Cells were kept in a stage top incubator (Tokai Hit) maintained at 37°C with 5% CO_2_ and protected from light.

### Fluorescent In Situ Hybridization

The RNA single-molecule FISH (smFISH) using 20mer DNA oligo probes was adapted from the work of Raj and colleagues^57^ and described in detail by Gaspar and colleagues^58^. In brief, DNA oligos were ordered from Integrated DNA. Technology and labeled in house with Cy3, Atto590, or Cy5. ACTB-MBS MEF cells were seeded on 12 mm glass coverslips (Electron Microscopy Service, 72290-03) that were coated for 30 minutes with 1:400 dilution of fibronectin, (Sigma-Aldrich F1141-2MG) in DPBS and cultured overnight. After fixation with 4% paraformaldehyde and permeabilization with 0.1% of Triton, cells were incubated with 20–40 nM probes in hybridization buffer for 3 hours at 37 °C. The unbound probes were washed away with 10% formamide and the coverslips were mounted on microscope slide using ProLong Diamond Antifade Mountant containing DAPI (Thermo Fisher Scientific, P36962) for nuclear staining.

FISH probes targeted the MBSv5 or MBSv1 region for U-2 OS cells or ACTB-MBS MEF cells, respectively. Internal controls *mGAPDH* and *mPolR2A* FISH probes targeted the ORF of each gene. Coverslips were mounted onto microscope slides with Prolong Diamond overnight and sealed with clear nail polish after curing the next day. The RNA FISH probe sequences are listed in **Supplementary Tables 2-6**.

For smFISH combined with immunofluorescence, 1:1000 dilution of rabbit anti-DCP1a (Abcam ab183709), 1:100 rabbit anti-XRN1 (Bethyl Laboratories A300-443A-M), 1:1000 rabbit anti-DDX6 (Bethyl Laboratories A300-461A), or 1:100 rabbit anti-G3BP (Aviva Systems Biology ARP37713_T100) were used as primary antibodies. A 1:5000 dilution of goat-anti-rabbit IgG (H+L) Alexa Fluor 750, Invitrogen A-21039, was used as the secondary antibody for all primary rabbit-derived antibodies.

### Fluorescence Microscopy

The fixed samples were imaged on an automated inverted Nikon Ti-2 wide-field microscope equipped with 60x, 1.4NA oil immersion objective lens (Nikon), Spectra X L.E.D. light engine (Lumencor), and Orca 4.0 v2 scMOS camera (Hamamatsu). The live cell experiments were performed on a custom microscope built around Nikon Ti-E stand. The excitation was through HTIRF (Nikon) with an LU-n4 four laser unit (Nikon) with solid state lasers with wavelengths 405, 488, 561, and 640 nm. The main dichroic was a quad band dichroic mirror (Chroma, ET-405/488/561/640 nm laser quad band set for TIRF applications). The imaging was done through the 100x 1.49NA oil immersion objective (Nikon). To achieve simultaneous 2-color imaging, we used a TriCam light splitter into three separate EMCCD cameras (Andor iXon Ultra 897) with ultraflat 2 mm thick imaging splitting dichroic mirrors (T565LPXR-UF2, T640LPXR-UF2). A band pass emission filter was placed in front of each camera, respectively (ET525/50 m, ET595/50 m, and ET655lp). The microscope was also equipped with an automated XY-stage with extra fine lead-screw pitch of 0.635 mm and 10 nm linear encoder resolution and a Piezo-Z stage (Applied Scientific Instrumentation) for fast Z-acquisition. A microscope stage top incubator (Tokai Hit, Model) is used to keep the sample at 37°C, 5% CO2 and saturating humidity. The whole microscope was under the control of Nikon Elements for automation.

### Live Cell Imaging

100,000 MEF cells stably expressing tet3G-FRB-SMG7C-IRES-FKBP-Halo-tdMCP and DDX6-eGFP were plated on a 35 mm with 20 mm micro well #1.5 cover glass bottom (Cellvis, D35-20-1.5-N). Cells were treated with 1 µg/mL Doxycycline overnight. The next morning, cells were incubated with 100nM JF646 Halo Ligand^59^ for 30 minutes, then rinsed once in complete DMEM with 10% FBS and 1% PenStrep, and transferred to a 37°C 5% CO_2_ incubator for at least 30 more minutes to equilibrate prior to imaging. During live-cell imaging, the cells were kept at 37°C with humidity control on a Tokai Hit stage top incubator. For data collection, each cell was imaged every 5 minutes for 2 hours with 100 ms exposure time. Low excitation conditions involved excitation with 2% 488 and 2% 640 laser stimulation, while the high excitation condition involved excitation with 2% 488 and 10% 640, stimulating light stress. Cells imaged after Sodium Arsenite stress were pre-treated with either 100 µM or 200 µM Sodium Arsenite 30 minutes prior to imaging in low excitation conditions. Analysis of live cell imaging was done using u-track v2^60^. P-bodies were detected in the eGFP-DDX6 channel using the Point Source Detection algorithm. The intensity of the FKBP-HaloTag-tdMCP channel was measured over time in the regions segmented by the detected P-bodies, and the intensity was summed at each time point. To produce the Supplementary Movies, max projected images were background subtracted in each channel using the rolling ball algorithm in ImageJ.

### Image analysis and quantification of RNA in smFISH experiments

We used an in-house RNA detection platform called uLocalize to count the compartmentalized P-body and cytoplasmic mRNAs separately. All custom code for smFISH-IF analysis and theoretical modeling can be found at (https://github.com/binwulab/uLocalize.git) and is summarized here:

To detect P-bodies in the DCP1a IF channel, we filter the image with Laplacian of Gaussian filter and segmented the area using an intensity threshold. Single RNAs were detected using a Local Maximum detection algorithm, and the single RNA intensity was determined by fitting the spot to a 3D Gaussian function to extract the center and the amplitude. To quantify RNAs in P-bodies, we measured the integrated intensity of the max projected RNA channel in the segmented P-body area. The integrated intensities of the RNA granules were normalized to RNA counts by dividing the median max projected single RNA intensity. Finally, RNA counts of all P-bodies in single cells were summed to obtain the total P-body RNA. counts used to fit the mathematical models. Cytoplasmic RNA was counted as all the detected RNA in the cytoplasm excluded from the segmented P-bodies. These RNA counts in cytoplasm and P-bodies were summed to obtain the total RNAs in single cells.

### Mathematical model to describe the compartmentalized RNA decay

Cytoplasmic and P-body RNA counts were calculated as described in Methods and fit to a mathematical model.

Model parameters definitions are described in **Figs. 3a-b**. The differential equations used to describe this simple kinetic model are as follows:

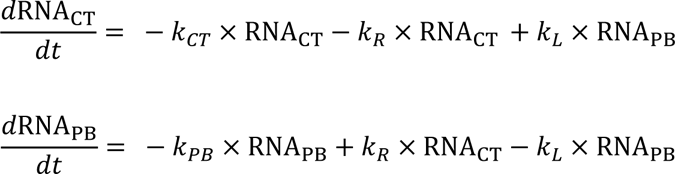

The differential equations were solved analytically in Mathematica. The solution was implemented in MATLAB to fit the cytoplasmic and P-body RNA counts simultaneously using the nonlinear Least Squares Fitting algorithm.

### Translation inhibition

25,000 ACTB-MBS MEF cells were plated on fibronectin-coated coverslips. Cells were treated with 1 µg/mL Doxycycline overnight to express the RIDR construct. The next morning, cells were incubated with 10nM JF503 Halo Ligand for 2 hours prior to fixation. Translation inhibitors were added to the sample 10 minutes prior to the Rapamycin + DRB or DRB control in each timepoint. Translation inhibitors were added at a final concentration of 100 µg/µL for Puromycin, 100 µg/µL for cycloheximide. smFISH-IF was then performed as described above.

### Western Blot

For DDX6 and XRN1 RNAi experiments, 100,000 ACTB-MBS MEF cells were treated as described in the Cell Culture and Transfection section. Cells were harvested by scraping and pelleted at 500xg for 2 minutes. Cell pellet was resuspended in 30 μl ice-cold lysis buffer (50mM HEPES pH 7.4, 150 mM KOAc, 15mM MgOAc2, 1% triton, leupeptin, pepstatin, PMSF, 1x EDTA-free Complete (Sigma 11873580001), 2 U Turbo DNase/ml (ThermoFisher AM2238), then gently pipetted 10 times to lyse. After incubating 5 minutes on ice, lysates were clarified by centrifugation at 20,000xg for 10 minutes at 4°C and the supernatant was transferred to a new tube. Samples were electrophoresed in a 4% SDS-PAGE gradient gel, transferred to a PVDF membrane, and blotted overnight with a 1:1000 dilution of rabbit anti-XRN1 primary antibody (Bethyl Laboratories A300-443A-M), 1:10000 dilution of rabbit anti-DDX6 primary antibody (Bethyl Laboratories A300-461A), or 1:250 rabbit anti-Ribosomal Protein S3 primary antibody (Santa Cruz sc-376008) as a control. Samples were then incubated with 1:5000 mouse anti-rabbit IgG HRP (Santa Cruz sc-2357) secondary antibody for 1 hour at room temperature then washed 3 times for 10 minutes in 1X Tris-Buffered Saline, 0.1% Tween® 20 Detergent (TBST). All incubation steps were done with gentle rocking. Samples were visualized on a Bio-Rad Chemidoc Imager.

## Code availability

The analysis code that supports the findings of this study is available in GitHub (https://github.com/binwulab/uLocalize.git)

## Acknowledgements

We thank members of the Wu and Inoue Lab for helpful discussion. We are also thankful to Boyang Hua for assistance with western blots. This work was supported by the National Institutes of Health (Grant # R01GM136897) and Pew Charitable Trust (Award ID 00030601) to B.W.. L.A.B. was supported by N.I.H. Training Grant (T32 GM008403).

## Author contributions

L.A.B. and B.W. designed the experiments. L.A.B. conducted all experiments and analysis. L.A.B. and B.W. wrote the paper. Y.L and T.I. advised on experimental design and interpreting data. B.W. supervised the project.

