## Supplementary material for "A Rapid Inducible RNA Decay system reveals fast mRNA decay in P-bodies": Supplemental Figures - Combined.pdf

#### Supplemental Figure 1

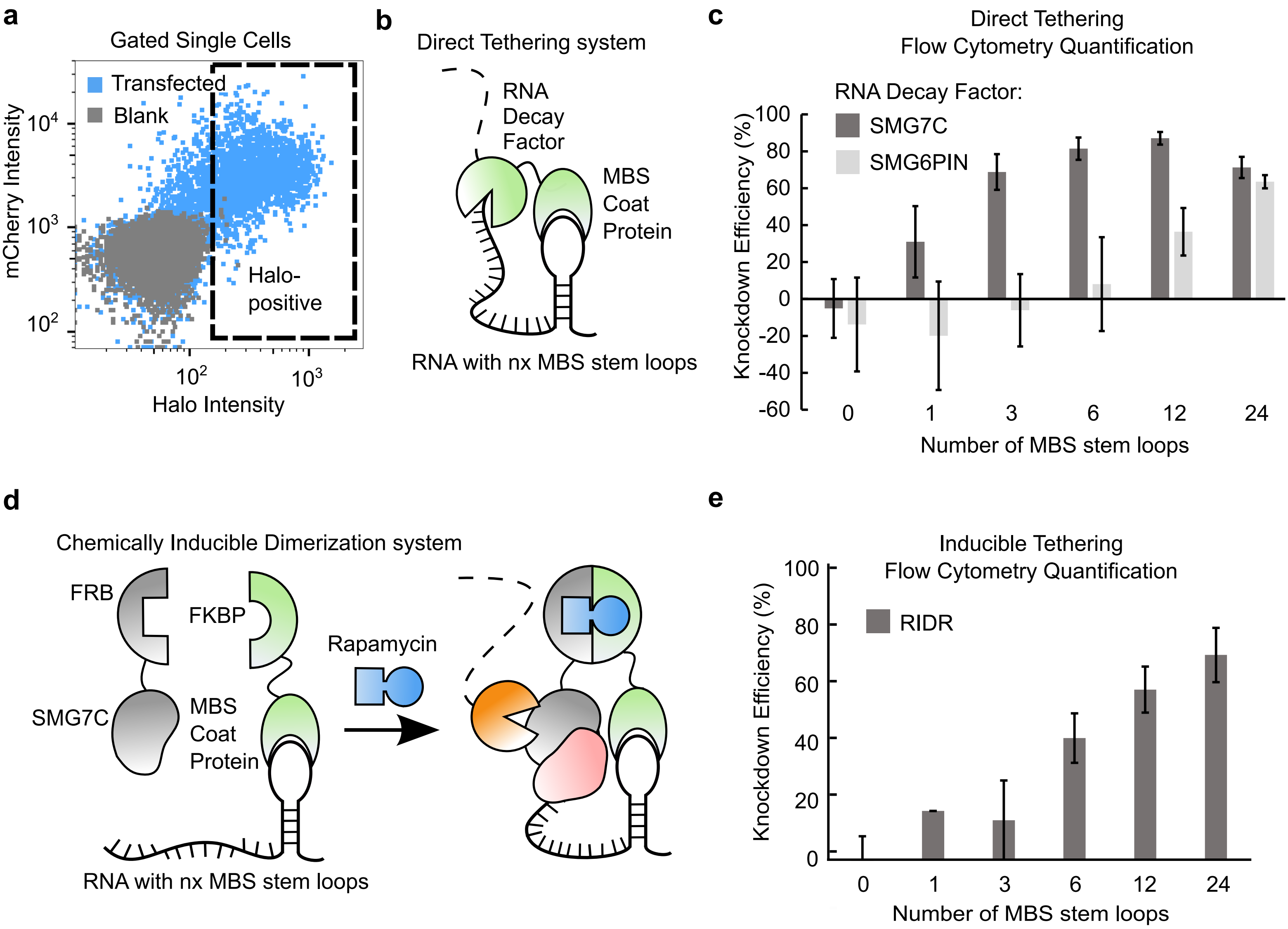

#### Supplemental Figure 2

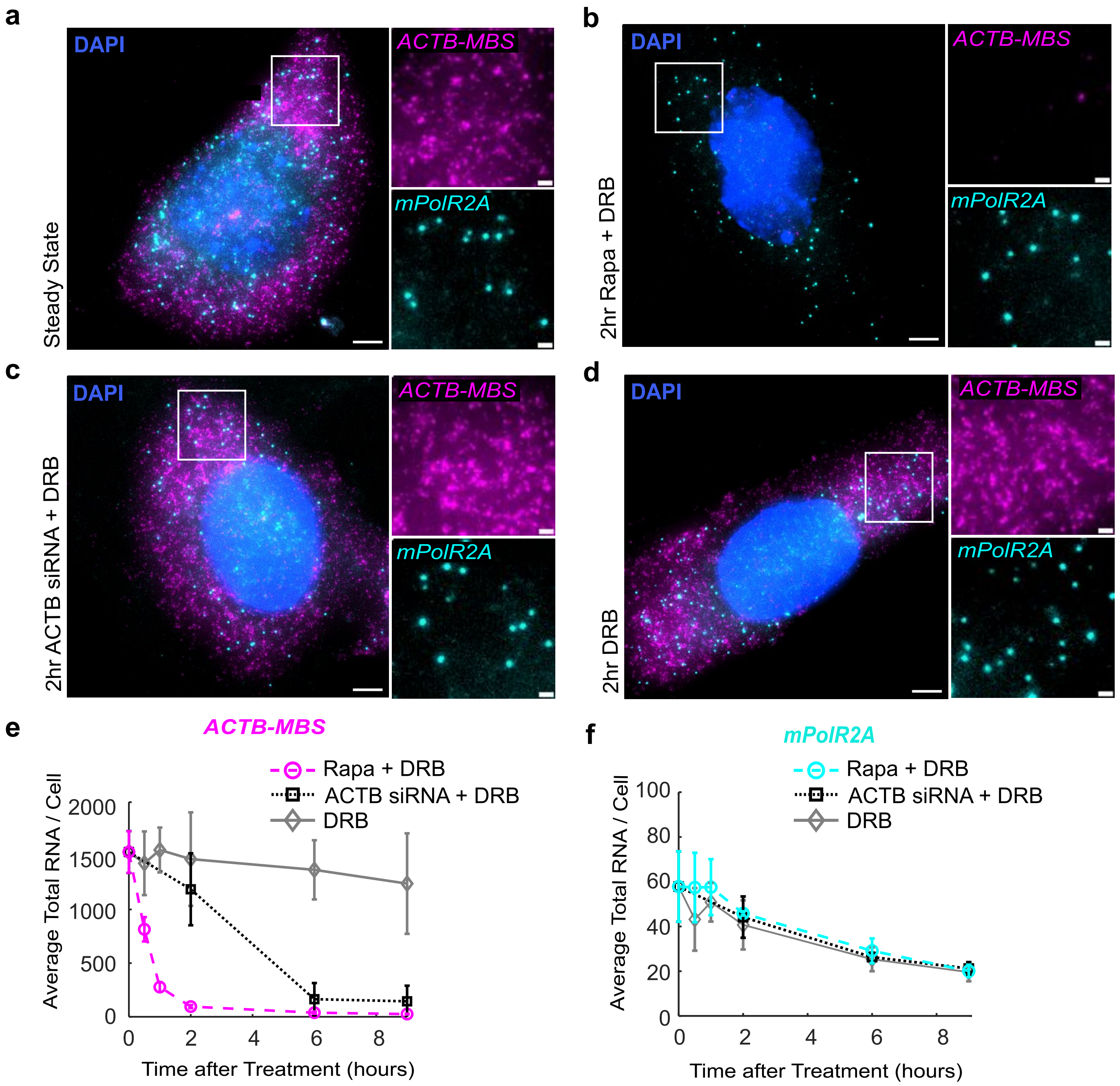

### Supplemental Figure 3

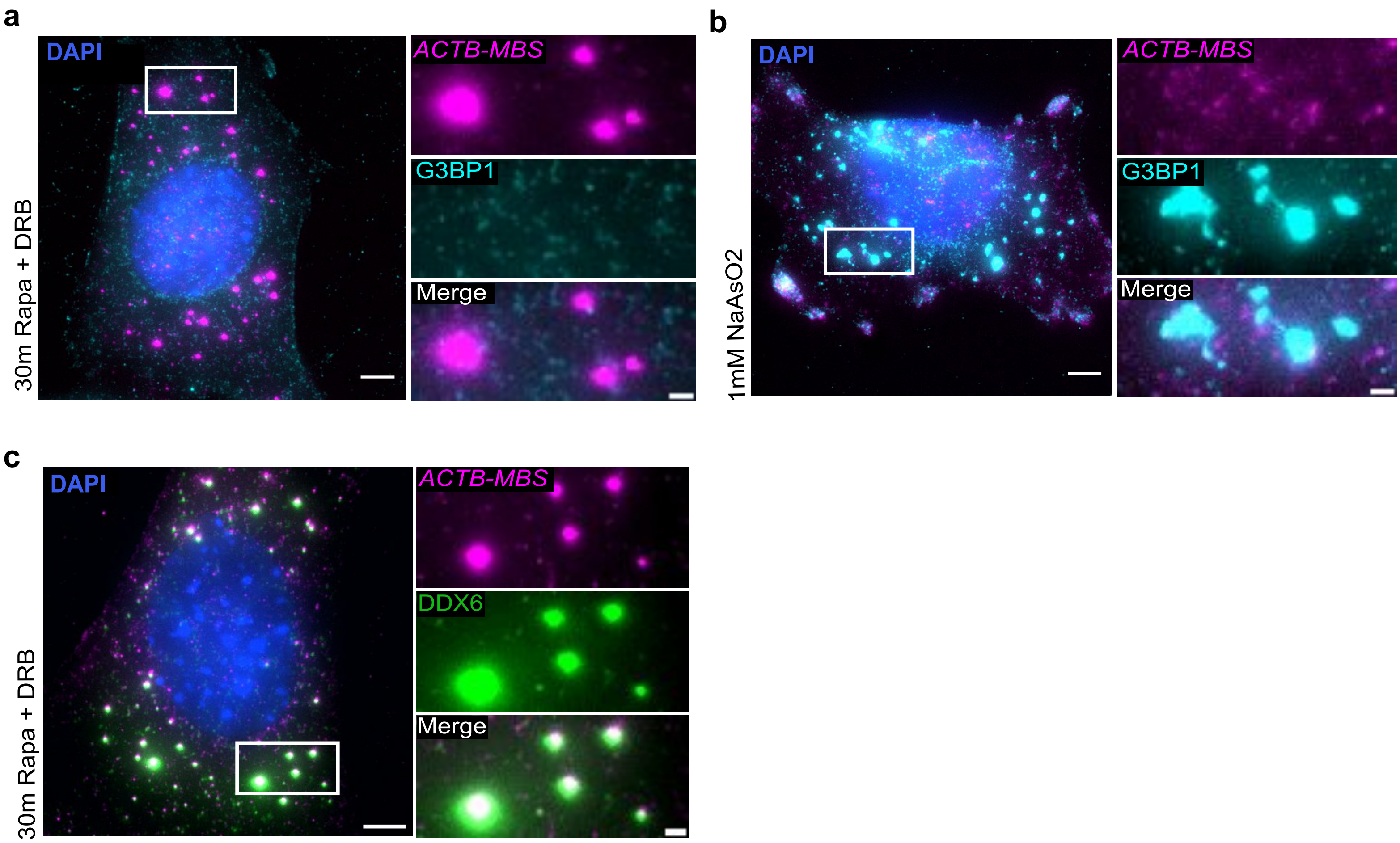

### Supplemental Figure 4

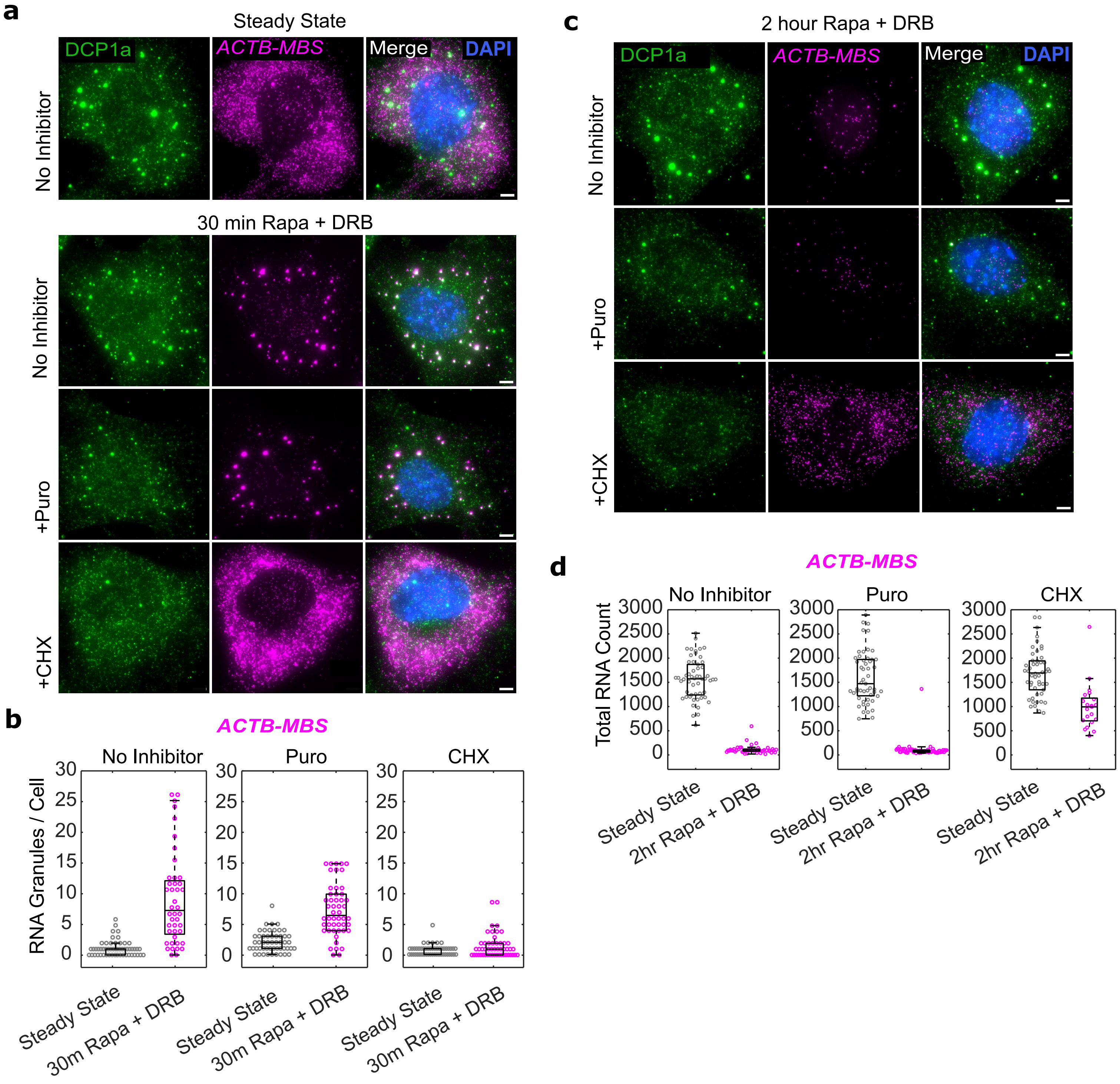

#### Supplemental Figure 5

**a**

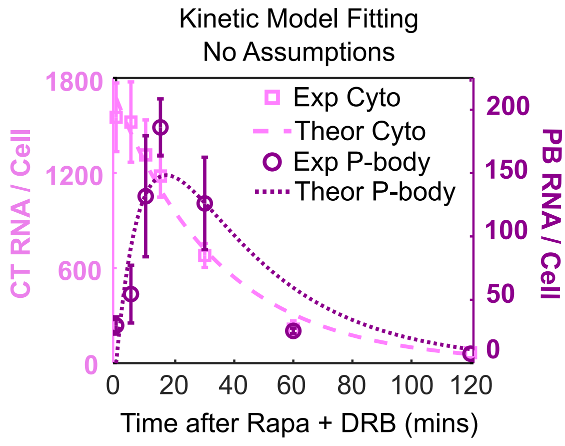


**b**

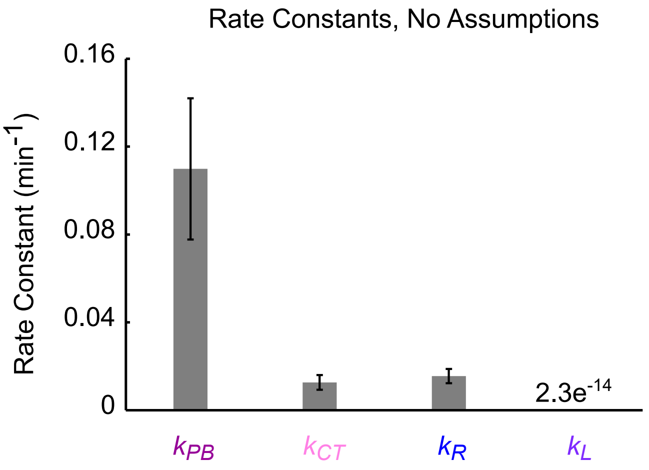

#### Supplemental Figure 6

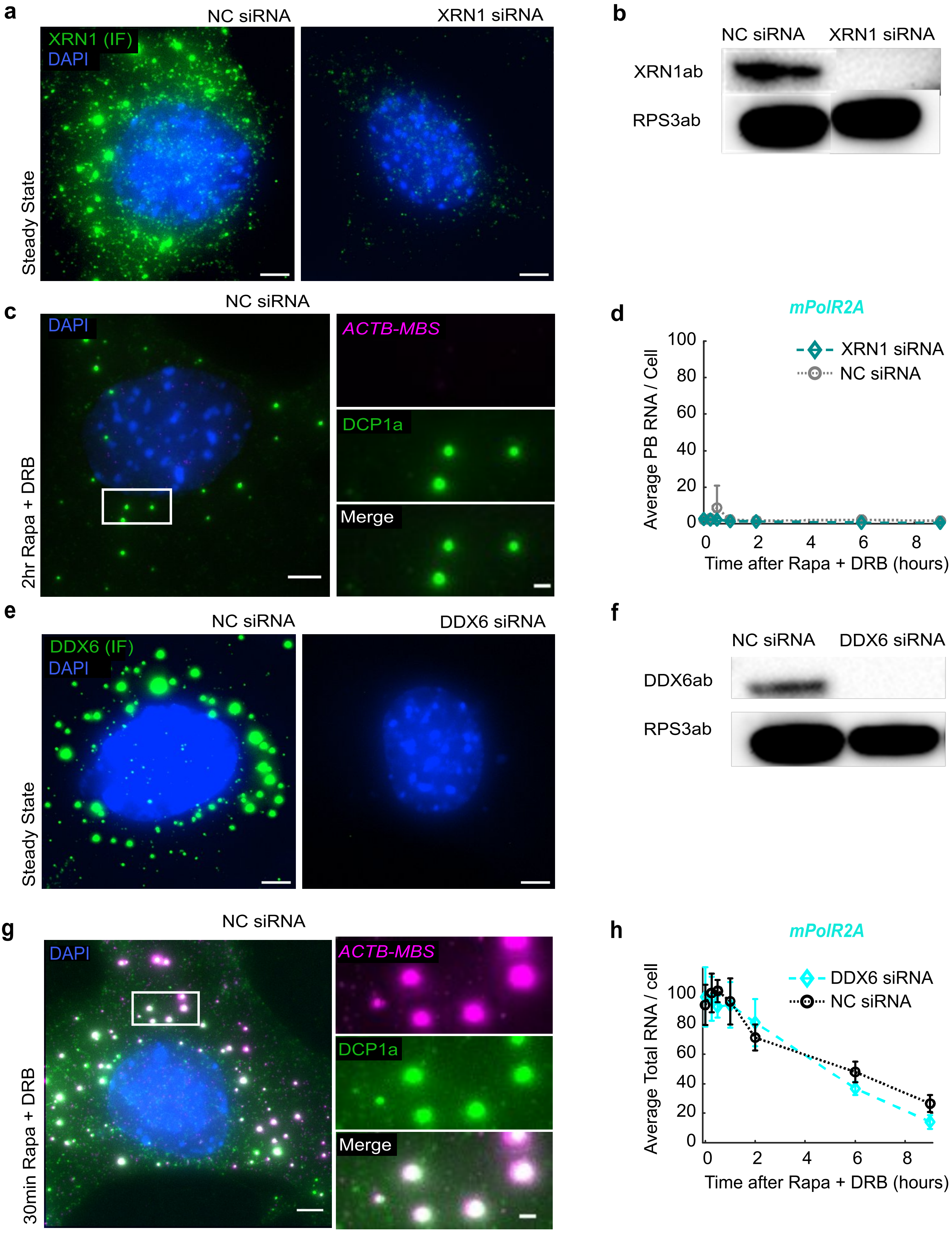
