## Supplementary material for "A Rapid Inducible RNA Decay system reveals fast mRNA decay in P-bodies": SupplementaryInformation.pdf

### Supplementary Figures

#### Supplementary Figure 1

Tethering SMG7C to 24xMBS produced highest knockdown efficiency.

#### Supplementary Figure 2

RIDR is fast, specific, and inducible on endogenous ACTB-MBS gene.

#### Supplementary Figure 3

Induced RNA granules colocalize with P-body marker DDX6 and not stress granule marker G3BP1.

#### Supplementary Figure 4

Translation inhibitors have different effects on RNA decay dynamics.

#### Supplementary Figure 5

Fitting to kinetic model with no assumptions shows similar results as Assumption III: $k_L$=0.

#### Supplementary Figure 6

Knockdown of DDX6 and XRN1 RNA is efficient.

### Supplementary Movies

#### Supplementary Movie 1

Live-cell imaging of ACTB-MBS MEF cells showing RNA granule formation.

#### Supplementary Movie 2

Live-cell imaging of ACTB-MBS MEF cells with high excitation conditions.

#### Supplementary Movie 3

Live-cell imaging of ACTB-MBS MEF cells with low excitation conditions.

#### Supplementary Movie 4

Live-cell imaging of ACTB-MBS MEF cells with low excitation conditions after 30-minute pre-treatment with 200µM Sodium Arsenite.

### Supplementary Tables

#### Supplementary Table 1

siRNA sequences ordered from IDT

#### Supplementary Table 2

MBSv1 20mer mixed smFISH probes

#### Supplementary Table 3

MBSv5 smFISH probes

#### Supplementary Table 4

hPolR2A smFISH probes

#### Supplementary Table 5

mPolR2A smFISH probes

#### Supplementary Table 6

mGAPDH smFISH probes

### Supplemental Figure Legends

**Supplemental Figure 1:** ***Tethering SMG7C to 24xMBS produced highest knockdown efficiency*** **a)** Gating strategy for all flow cytometry experiments demonstrating how single HEK293T cells were gated. Representative flow data of cells that were co-transfected with mCherry-nxMBS and fusions of HaloTag-tdMCP with an RNA decay factor or CID protein (blue) or nothing (gray). HaloTag-positive cells (dashed black box) were selected for the quantification of mCherry knockdown for each condition relative to a negative control. **b)** Schematic of the direct tethering of a general RNA decay factor to an MBS by fusing the RNA decay factor to the tandem MS2 coat protein (tdMCP) **c)** HEK293T cells were transiently transfected with mCherry-nxMBS (where n = 0, 1, 3, 6, 12, 24) and SMG7C-HaloTag-tdMCP or SMG6PIN-HaloTag-tdMCP constructs. HaloTag was labelled with JF503 Halo-ligand. The fluorescence of single cells was measured by flow cytometry. The knockdown efficiency was calculated for each condition with respect to a negative control with no decay factor, Halo-tdMCP. **d)** Schematic of the inducible mRNA decay system using FRB/FKBP CID induced by rapamycin ligand. SMG7C-fused FRB is tethered to target RNA with MBS by FKBP-tdMCP to cause its degradation. **e)** Knockdown efficiencies of mCherry-nxMBS were quantified from flow cytometry experiments using the inducible system shown in (c). The knockdown efficiency is calculated for each condition with respect to itself when Rapa is not added. 1000-3000 HaloTag-positive cells were quantified per condition. Error bars represent the standard deviation across 2 biological replicates.

**Supplemental Figure 2:** ***RIDR is fast, specific, and inducible on endogenous ACTB-MBS gene.*** **a-d)** Representative smFISH images of ACTB-MBS MEF cells stably expressing RIDR in steady state conditions **(a)**, after 2 hours Rapa + DRB treatment **(b)**, after 2 hours ACTB siRNA treatment **(c)**, and after 2 hours of DRB treatment alone **(d)**. The white boxes were enlarged on the right. *ACTB-MBS* FISH: magenta; *mPolR2A* FISH: cyan; DAPI: blue. Scale bars: 5 µm for original and. 1 µm for zoomed images. **e-f)** Quantification of time-resolved two-color smFISH experiment over 9-hours after induction with different treatments. The number of transcripts for *ACTB-MBS* **(e)** and *mPolR2A* **(f)** were counted in the same cells for all time points. Rapa + DRB: circles; ACTB siRNA + DRB: squares; DRB alone: diamonds. Error bars represent standard deviation of the means of 4 replicates (150-284 cells were quantified per condition)

**Supplemental Figure 3:** ***Induced RNA granules colocalize with P-body marker DDX6 and not stress granule marker G3BP1*** smFISH-IF experiments were conducted on ACTB-MBS MEF cells stably expressing RIDR construct to ascertain the identity of the observed RNA granules. *ACTB-MBS* FISH: magenta; SG marker G3BP1 IF: cyan; P-body marker DDX6 IF: green; DAPI: blue. **a**) After 30 minutes Rapa induction, there is ACTB-MBS mRNA granule, but no stress granule formation. **b**) The G3BP1 antibody label the stress granule after cells were treated with 1 mM Sodium Arsenite for 10 minutes, demonstrating the efficacy of G3BP1 antibody. **c)** After 30 minutes induction, the induced RNA granules colocalize with another P-body marker DDX6. White boxes were enlarged on the right for each. Scale bars: 5 µm for original images, 1 µm for zoomed images.

**Supplemental Figure 4: *Translation inhibitors have different effects on RNA decay dynamics.*** Different Translation inhibitors are combined with RIDR to study effect on RNA decay behavior **a)** Representative smFISH-IF images of ACTB-MBS MEF cells expressing RIDR construct then treated with no inhibitor, puromycin, or cycloheximide for 30 mins, followed by induction by rapamycin plus DRB for 30 min **b)** Quantification of *ACTB-MBS* RNA granule counts in Steady State vs 30 min Rapa + DRB after no translation inhibition, or treatment with puromycin or cycloheximide. **c)** Representative FISH-IF images of ACTB-MBS MEF cells expressing RIDR construct then treated with no inhibitor, puromycin, or cycloheximide for 30 mins, followed by induction by rapamycin plus DRB for 2 hours **d)** Quantification of (a) showing *ACTB-MBS* RNA counts in Steady State vs 2-hour Rapa + DRB after no translation inhibition, or treatment with puromycin or cycloheximide. 150-284 cells were quantified per condition. Scale bars: 5 µm.

**Supplementary Figure 5: *Fitting to kinetic model with no assumptions shows similar results as Assumption III: $k_L$=0*** **a)** Fitting results with no assumptions. RNA counts in the P-body: dark magenta; RNA counts in the cytoplasm: light magenta; Experimental data: symbols; Theoretical fit: lines. Error bars represent standard deviation of the means of 4 replicates. **b)** Model parameters determined from fitting with no assumptions. Error bars indicate the standard deviation of the fitted parameters across 4 replicates.

**Supplemental Figure 6. *Knockdown of DDX6 and XRN1 RNA is efficient*.** ACT-MBS MEF cells were treated with siRNA against XRN1 (**a-d),** DDX6 (**e-h**) or scrambled siRNA (NC) **(a-h**) for 72 hours. Cells were treated with Rapa and DRB, and fixed at different time points. **a)** Representative IF images for cells treated with NC (left) or XRN1 (right) siRNA. XRN1 IF: green; DAPI: blue. **b)** Western blots with XRN1 antibody on cells treated with NC (left column) or XRN1 (right column) siRNA with RPS3 antibody as an internal control. **c)** Representative smFISH-IF image for XRN1 siRNA treated cell at 2 hours post-induction. The white box was enlarged on the right. *ACTB-MBS* FISH: magenta; DCP1a IF: green; DAPI: blue. **d)** Quantification of mPolR2A mRNAs in P-bodies over 9-hour time course after induction. XRN1 siRNA: cyan; NC siRNA: gray. **e)** Representative IF images for cells treated with NC (left) or DDX6 (right) siRNA. DDX6 IF: green; DAPI: blue. **f)** Western blots with DDX6 antibody on cells treated with NC (left column) or DDX6 (right column) siRNA with RPS3 antibody as an internal control. **g)** Representative smFISH-IF image for DDX6 siRNA treated cell at 30 minutes post-induction. The white box was enlarged on the right. *ACTB-MBS* FISH: magenta; DCP1a IF: green; DAPI: blue. **h)** Quantification of total *mPolR2A* mRNAs per cell over 9-hour time course after induction. DDX6 siRNA: cyan; NC siRNA: black. Error bars represent the standard deviation of the means of 4 replicates (203-450 cells were quantified per condition). Scale bars: 5 µm for original images, 1 µm for zoomed images.

### Supplementary Movie Legends

**Supplementary Movie 1. *Live-cell imaging of ACTB-MBS MEF cells showing RNA granule formation.*** Live cell imaging was done over 2 hours to visualize *ACTB-MBS* (FKBP-HaloTag-tdMCP, magenta). This cell was imaged at 5-minute time intervals using low excitation laser power. *ACTB-MBS* RNA granules can be observed within 10 minutes and disappear within 1 hour. Scale bar: 5 µm.

**Supplementary Movie 2. *Live-cell imaging of ACTB-MBS MEF cells with high excitation conditions.*** Live cell imaging was performed to track P-bodies (DDX6-eGFP, green, left) and *ACTB-MBS* (FKBP-HaloTag-tdMCP, magenta, middle) to assess their colocalization (merged image, right) after induction. This is the full movie of the montages displayed in **Fig. 5a** showing a representative ACTB-MBS MEF cells treated with Rapa then imaged at 5-minute time intervals with high excitation laser power. Scale bar: 5 µm.

**Supplementary Movie 3. *Live-cell imaging of ACTB-MBS MEF cells with low excitation conditions.*** Live cell imaging was performed to track P-bodies (DDX6-eGFP, green, left) and *ACTB-MBS* (FKBP-HaloTag-tdMCP, magenta, middle) to assess their colocalization (merged image, right) after induction. This is the full movie of the montages displayed in **Fig. 5b** showing a representative ACTB-MBS MEF cells treated with Rapa then imaged at 5-minute time intervals with the minimal laser power sufficient to observe RNA granules colocalized with P-bodies. Scale bar: 5 µm.

**Supplementary Movie 4. *Live-cell imaging of ACTB-MBS MEF cells with low excitation conditions after 30-minute pre-treatment with 200µM Sodium Arsenite.*** Live cell imaging was performed to track P-bodies (DDX6-eGFP, green, left) and *ACTB-MBS* (FKBP-HaloTag-tdMCP, magenta, middle) to assess their colocalization (merged image, right) after induction. This is the full movie of the montages displayed in **Fig. 5c** showing a representative ACTB-MBS MEF cells treated with Rapa then imaged at 5-min time intervals with minimal laser power after cells were pretreated with 200 µM Sodium Arsenite for 30 minutes. Scale bar: 5 µm.
